# ITPase Deficiency Causes Martsolf Syndrome With a Lethal Infantile Dilated Cardiomyopathy

**DOI:** 10.1101/383612

**Authors:** Mark T. Handley, Kaalak Reddy, Jimi Wills, Elisabeth Rosser, Archith Kamath, Mihail Halachev, Gavin Falkous, Denise Williams, Phillip Cox, Alison Meynert, Eleanor S. Raymond, Harris Morrison, Stephen Brown, Emma Allan, Irene Aligianis, Andrew P Jackson, Bernard H Ramsahoye, Alex von Kriegsheim, Robert W. Taylor, Andrew J. Finch, David R. FitzPatrick

**Author notes:** These authors contributed equally to this work (Mark T. Handley, Kaalak Reddy). Address for correspondence: David R. FitzPatrick.

## Abstract

Martsolf syndrome is characterized by congenital cataracts, postnatal microcephaly, developmental delay, hypotonia, short stature and biallelic hypomorphic mutations in either *RAB3GAP1* or *RAB3GAP2.* Through genetic analysis of 85 unrelated “mutation negative” probands referred with Martsolf syndrome we identified two individuals with different homozygous null mutations in *ITPA*, the gene encoding inosine triphosphate pyrophosphatase (ITPase). The probands reported here each presented with a lethal and highly distinctive disorder; Martsolf syndrome with infantile-onset dilated cardiomyopathy. Severe ITPase-deficiency has been previously reported with infantile epileptic encephalopathy (MIM 616647). ITPase acts to prevent incorporation of inosine bases (rl/dl) into RNA and DNA. In *Itpa*-null cells, dI was undetectable in genomic DNA. dI could be identified at a low level in mtDNA but this was not associated with detectable mitochondrial genome instability, mtDNA depletion or biochemical dysfunction of the mitochondria. rI accumulation was detectable in lymphoblastoid RNA from an affected individual. In *Itpa*-null mouse embryos rI was detectable in the brain and kidney with the highest level seen in the embryonic heart (rI at 1 in 385 bases). Transcriptome and proteome analysis in mutant cells revealed no major differences with controls. The rate of transcription and the total amount of cellular RNA also appeared normal. rI accumulation in RNA – and by implication rI production - correlates with the severity of organ dysfunction in ITPase deficiency but the basis of the cellulopathy remains cryptic. While we cannot exclude cumulative minor effects, there are no major anomalies in the production, processing, stability and/or translation of mRNA.

**Author Summary:** Nucleotide triphosphate bases containing inosine, ITP and dITP, are continually produced within the cell as a consequence of various essential biosynthetic reactions. The enzyme inosine triphosphate pyrophosphatase (ITPase) scavenges ITP and dITP to prevent their incorporation into RNA and DNA. Here we describe two unrelated families with complete loss of ITPase function as a consequence of disruptive mutations affecting both alleles of *ITPA*, the gene that encodes this protein. Both of the families have a very distinctive and severe combination of clinical problems, most notably a failure of heart muscle that was lethal in infancy or early childhood. They also have features of another rare genetic disorder affecting the brain and the eyes called Martsolf syndrome. We could not detect any evidence of dITP accumulation in genomic DNA from the affected individuals. A low but detectable level of inosine was present in the mitochondrial DNA but this did not have any obvious detrimental effect. The inosine accumulation in RNA was detectable in the patient cells. We made both cellular and animal models that were completely deficient in ITPase. Using these reagents we could show that the highest level of inosine accumulation into RNA was seen in the embryonic mouse heart. In this tissue more than 1 in 400 bases in all RNA in the cell was inosine. In normal tissues inosine is almost undetectable using very sensitive assays. The inosine accumulation did not seem to be having a global effect on the balance of RNA molecules or proteins.

## Introduction

It is 40 years since two brothers were reported with severely delayed neurocognitive development, spasticity, postnatal microcephaly, short stature, congenital cataracts and primary hypogonadism[1], characterising a disorder that is now termed Martsolf syndrome (MIM 212720). Warburg Micro syndrome (MIM 600118, 614225, 615222, 615663) is an overlapping condition that was described in 1993, which also has microphthalmia/microcornea, retinal dystrophy, optic nerve atrophy and intracranial malformations as clinical features[2]. 60% of cases referred to us with a diagnosis of Warburg Micro syndrome have loss-of-function mutations in either *RAB3GAP1, RAB3GAP2, RAB18* or TBC1D20[3-6]. 44% of Martsolf syndrome cases have mutations in *RAB3GAP1* or *RAB3GAP2*, which perturb but do not completely abolish the expression or function of the encoded protein%[7, 8]. The relatively high proportion of unexplained cases in both syndromes indicates that there are likely to be more disease loci and/or causative genetic mechanisms to be discovered.

Infantile-onset dilated cardiomyopathy (iDCM) is a rare, aetiologically heterogeneous disorder that may present as acute, commonly lethal, and with cardiogenic shock [9]. Isolated iDCM may be caused by genetically determined primary abnormalities of heart muscle (sarcomere, Z-disc, desmosomes etc) while iDCM as a component of a multisystem disorder is most commonly secondary to an inborn error of metabolism (**Table 1**)[10-12] with the prognosis being dependent on the underlying cause. Early genetic testing is recommended in iDCM as it may help direct the clinical management[13].

**Table 1:** Genetic causes of syndromic infantile-onset dilated cardiomyopathy.

|  | OMIM | Genes | Progressive Neurological Disorder | Eye Disease | Other Features |
| --- | --- | --- | --- | --- | --- |
| e | #203800 | *ALMS1* | No | Yes | AR. Systemic fibrosis, multiple organ involvement, retinal degeneration, hearing loss, childhood obesity, diabetes mellitus, urogenital dysfunction. Pulmonary, hepatic and renal failure. |
|  | #302060 | *TAZ* | No | No | X-linked recessive. Neutropenia, growth delay and skeletal myopathy. Gross motor delay, early death from septicaemia. Intermittent lactic acidemia, muscle weakness. Endocardial fibroelastosis. |
| (3-luria | #610198 | *DNAJC19* | +/- | No | AR. Prenatal or postnatal growth failure, cerebellar ataxia, significant motor delay, testicular dysgenesis-cryptorchidism, severe perineal hypospadias, elevated plasma and urine 3-methylglutac acid and 3-methylglutaric acid. |
| e II | #600649 | *CPT2* | No | No | AR. Recurrent hypoglycaemia, seizures, liver failure and transient hepatomegaly. Episodes triggered by infections, fever, or fasting. Elevated creatine kinase. |
| e | #605676 | *DSP* | No | No | AR. Cardiocutaneous syndrome: woolly hair, palmoplantar keratoderma, tooth agenesis. |
| ated | #613642 | *SDHA* | No | No | Normal growth and neuromuscular examination. Normal development, no seizures. Normal brain MRI. |
| oxyl<br>se | #609016 | *HADHA* | Yes | Yes | AR. Coma, hypoglycaemia, myopathy, hepatopathy, intellectual disability, infantile spasms (-/+). Brain imaging: atrophic changes to parieto-occipital lesions, abnormal EEG background. Choro-retinal changes, abnormal ERG. |
| e | #212140 | *SLC22A5* | Yes | No | AR. Hypoketotic hypoglycaemia (encephalopathy), hepatomegaly, muscle weakness, cardiac arrhythmia, asymptomatic to sudden death. |

Here we report two families with a very distinctive clinical presentation of lethal iDCM and Martsolf syndrome associated with homozygous null mutations in *ITPA* which encodes inosine triphosphate pyrophosphatase (ITPase). ITPase is an enzyme that functions to prevent incorporation of inosine bases (rI/dI) into RNA and DNA by scavenging ITP/dITP in the cell. An autosomal recessive partial deficiency of inosine triphosphate pyrophosphatase (ITPase) has been recognised since the late 1960’s via accumulation of inosine triphosphate (ITP) in erythrocytes[14]. This is a relatively common trait that is clinically asymptomatic although it may influence sensitivity to certain drugs[15]. The trait is caused by hypomorphic mutations in *ITPA* (the gene encoding ITPase) which affect splicing and/or protein stability[16]. Biallelic loss-of-function mutations in *ITPA* have recently been reported as the cause of an early infantile encephalopathy (EIEE35, MIM #616647)[17]. We present data testing and refuting various hypotheses regarding the molecular consequences of ITPase deficiency on the genome, transcriptome and proteome.

## Results

### Clinical Information

In Family 4911 (**Figure 1A**) a maternal uncle (4911 V:5) and aunt (4911 V:7) of the proband (VI:3) had been described in a clinical paper as Martsolf syndrome with a previously unreported association with an early-onset cardiomyopathy[18]. The proband in the present study, their nephew, died at the age of 2 years. No post mortem examination was carried out and the exact cause of his death could not be confirmed. Prior to his demise he had been noted to have postnatal onset microcephaly with severe delay in all aspects of his development. He had bilateral cataracts diagnosed at the age of 13 months. Generalised seizures began at the age of 14 months. He was noted to have small genitalia. A clinical diagnosis of Martsolf syndrome was made and he had a negative screen for *RAB3GAP1, RAB3GAP2, RAB18* and *TBC1D20.* His elder brother, two maternal uncles and a maternal aunt all had a very similar pattern of problems (**Table 2**) and all had died in early childhood with evidence of cardiac failure[18]. Serial echocardiograms in 4911 VI:3 had shown persistent but mild dilation of his left ventricle and he is assumed to have died as a result of the progression of his cardiac disease.

**Figure 1.**
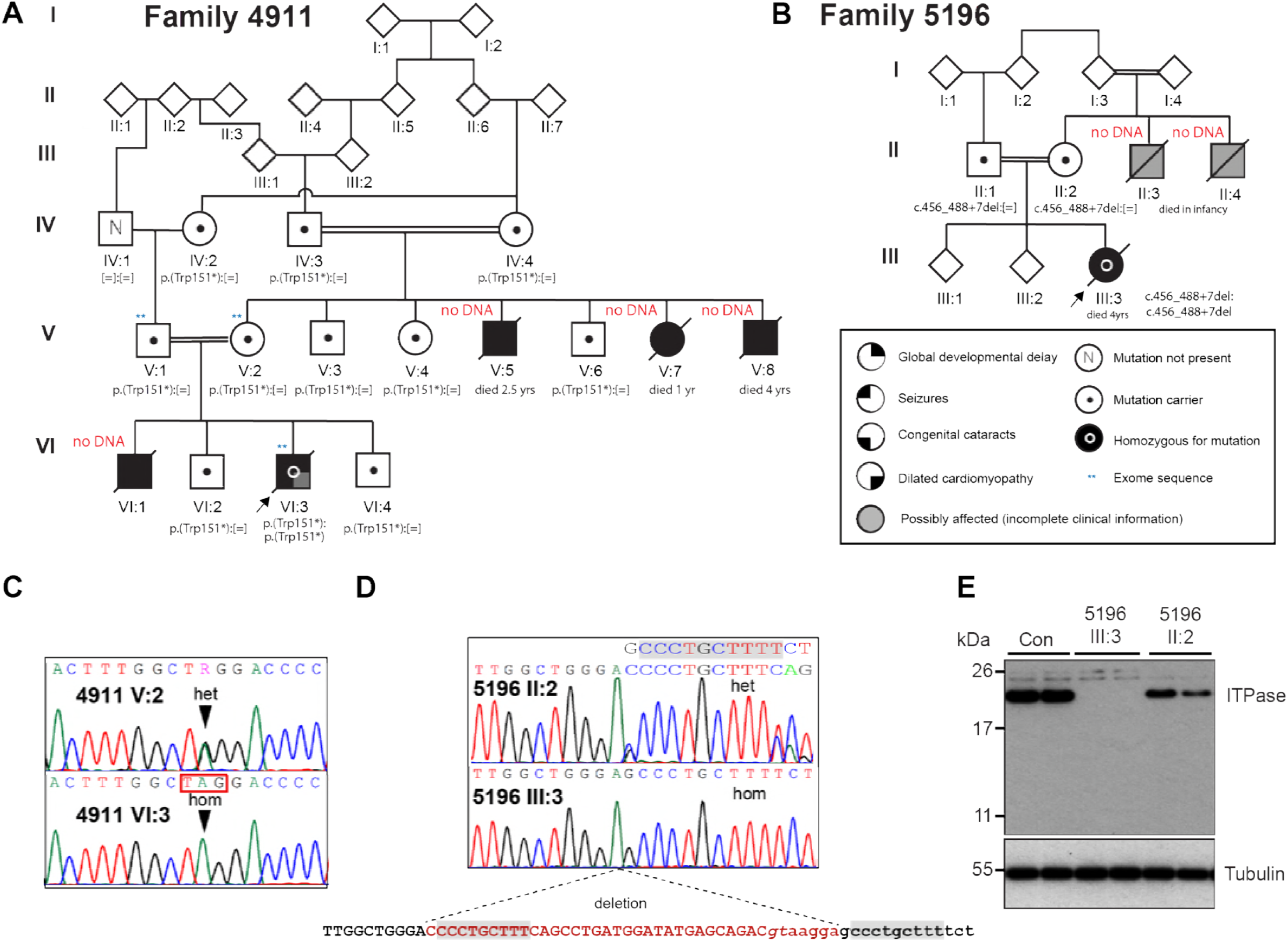
Loss-of-function mutations in *ITPA* identified in Martsolf syndrome with infantile cardiomyopathy. (A) Pedigree for Family 4911 showing the transmission of the *ITPA* c.452G>A, p. Trp151* allele. (B) Pedigree for Family 5196 showing the transmission of the *ITPA* c.456_488+7del allele. Electropherograms from sequencing of the affected individual (lower) and his mother (upper) from Family 4911 (C) and the affected individual (lower) and her mother (upper) from Family 5196 (D) show the sequence changes. The mutation nomenclature is based on the reference sequences NM_033453 and NP_258412. (E) Western blotting shows that ITPA protein is absent in a lymphoblastoid cell line derived from the affected individual 5196 III:3 and reduced in a line derived from her mother. Blotting for Tubulin serves as a loading control and each lane on the blot corresponds to an individual lysate sample.

**Table 2:** Clinical Features.

|  | Family 4911 |  |  |  |  | Family 5196 |
| --- | --- | --- | --- | --- | --- | --- |
|  | V:5 | V:7 | V:8 | VI:1 | VI:3 | III:3 |
| Ethnicity | Pakistan |  |  |  |  | Pakistan |
| Consanguinity | Yes |  |  |  |  | Yes |
| Family history | See Pedigree Figure 1A |  |  |  |  | See Pedigree Figure 1B |
| Sex | Male | Female | Male | Male | Male | Female |
| Age at last assesment | 9 mo | 9 mo |  | not seen | 15 months | 4 yrs |
| Age at death | 2.5 years | 1yr | 4yr | 6 months | 2yrs | 4yrs |
| **Birth and newborn** |  |  |  |  |  |  |
| Gestation | 40 | 40 | 37 | ? | 40 | 39 |
| Birth weight g [z score] | 3200 [-0.71] | 2800 | 2070 | ? | ? | 2892 |
| **Developmental milestones** |  |  |  |  |  |  |
| Social smile |  | >9 months | ? | ? | ? | 8 months |
| Sat Independently |  |  |  |  |  | Not achieved |
| Walked Independently | Not achieved | Not achieved | Not achieved | Not applicable | Not achieved | Not achieved |
| First words | Not achieved | Not achieved | Not achieved | Not applicable | Not achieved | Not achieved |
| **Growth parameters** |  |  |  |  |  |  |
| Height/length cm [z score] | 71 [-0.33] | 67 [-1.3] | ? | ? | ? | 97 [-1.1] |
| Weight kg [z score] | 7.7 [-1.7] | 6.5 [2.7] | ? | ? | ? | 10.2 [-4.2] |
| OFC cm [z score] | 40 [-5.4] | 39.7 [-4.8] | ? | ? | ? | 42 [-7.6] |
| **Clinical features** |  |  |  |  |  |  |
| Neurology | Hypotonic, brisk reflexes | Hypotonic, brisk reflexes, poor bulbar function | Hypotonia. OFC< 3rd centile. Optic atrophy | Delayed | Hypotonia. Developmental delay | Hypotonia, Severe developmental delay, microcephaly |
| Seizures | None | Started 8 months, facial twitching |  | None | started 14 months, generalised | Presented at 5 months with status epilepticus |
| EEG |  | Multifocal spikes and sharp waves | bursts of irregular activity |  | normal | normal |
| Neuroimaging | CT Brain showed generalised cerebral atrophy | CT and MRI of brain; generalised cerebral and brainstem atrophy | None | None | None | Neuropathology: see Figure 2 |
| Cardiac | Cardiac failure | Dilated cardiomyopathy | Dilated cardiomyopathy | Presented in cardiac failure at 6 months of age. Died suddenly before formal assessment | Mild dilation of left ventricle | See Figure 2 |
| Eye/vision | Central lens opacities | Bilateral Cataracts | Bilateral Cataracts | Bilateral cataracts. Developed between 2 and 6 months of age | Bilateral cataracts detected 13 months | Bilateral cataracts |
| Additional malformations | Cryptorchidism, slight hirsutism | Slight hirsutism | Microcephaly. Cryptorchidism |  | Small genitalia, microcephaly | Severe thymic atrophy, gracile bones on skeletal survey |
| ITPA variants (GRCh37/hg19) | No DNA available | No DNA available | No DNA available | No DNA available | chr20<br>g.3202527G>A;<br>c.452G>A,<br>p.Trp151*<br>(rs200086262);<br>homozygous | chr20 hg19<br>g.3202531-3202570del;<br>c.456_488+7del;<br>homozygous |

In family 5196 (**Figure 1B**) the affected proband (5196 III:3) was a girl who died at the age of 4 years. She had previously been diagnosed as having Martsolf syndrome on the basis of profound developmental delay, failure to thrive, microcephaly, seizures and congenital cataracts. Screening of the known Martsolf syndrome and Warburg Micro genes was negative. She presented in severe cardiac failure and died shortly after this. She had not previously been suspected of having any cardiac disease. In addition to the known anomalies, a post mortem examination revealed marked dilation of the left ventricle with increased trabeculation and mild fibroelastosis (**Figure 2A-C**). Fatty infiltration was noted of the right ventricle (**Figure 2B**). Neuropathology showed cerebellar atrophy (**Figure 2F**), microgliosis of dentate and olivary nuclei, vacuolation of white matter (**Figure 2E**) with scattered axonal spheroids (**Figure 2H**) and gliosis of the hippocampus. 5196 III:3 had two maternal uncles who died in infancy (5196 II:3 and 5196 II:4; **Figure 1B**) who were suspected of having the same disorder although no clinical details were available from either individual.

**Figure 2:**
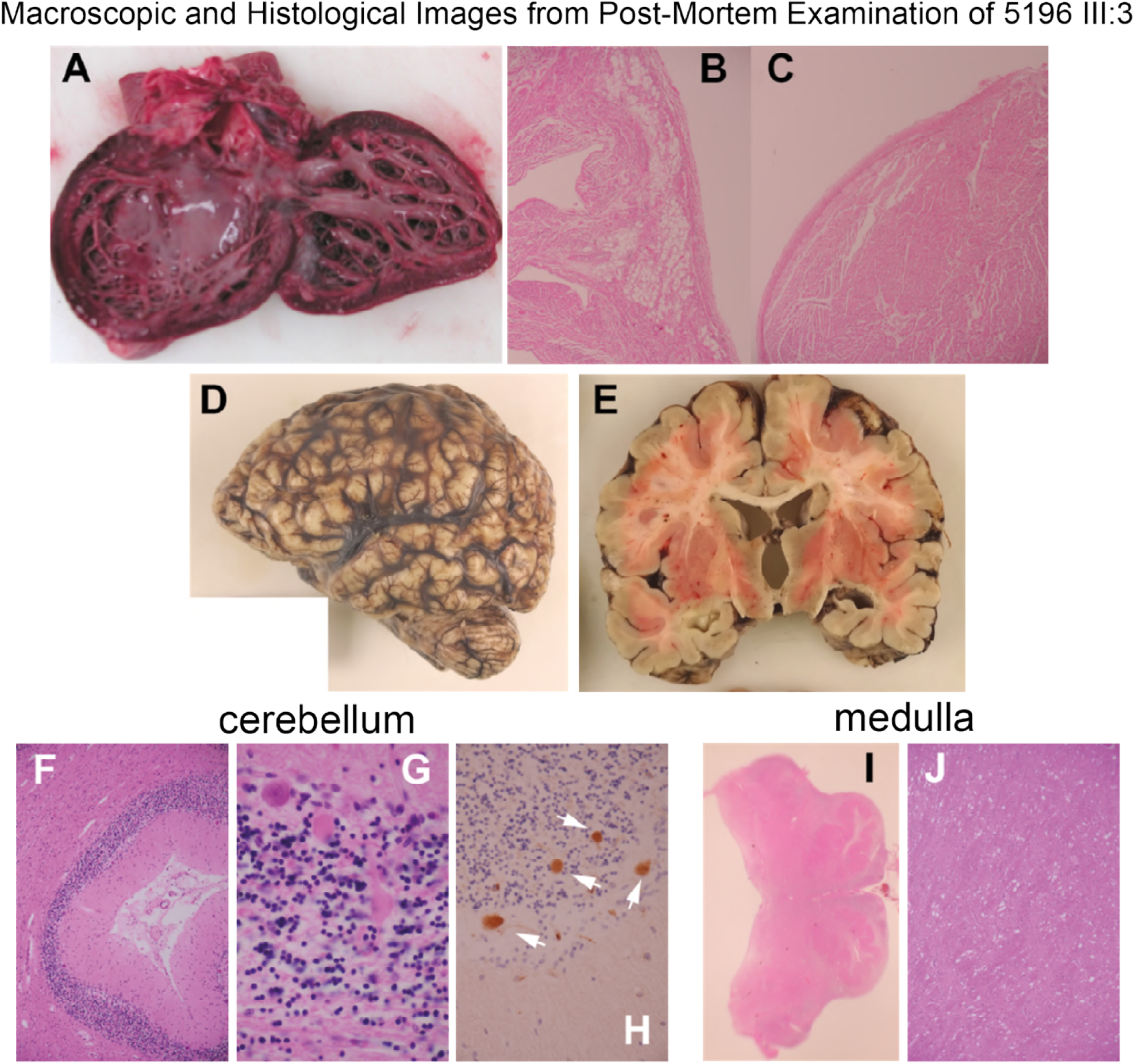
Phenotype associated with complete loss of ITPase function. (A) Photograph of the dissected heart from the affected individual (III:3) in Family 5196 showing increased trabeculation and fibroelastosis of the endocardium. (B,C) Photomicrographs (x200) showing fatty infiltration and fibroelastosis of the heart respectively (D) Photograph of the fixed brain showing microcephaly/micrencephaly. (E) Coronal slice through brain showing dilatation of the lateral and third ventricles with reduced volume of white matter. (F) Image (x100) showing atrophy of the cerebellar cortex. (G) Photomicrograph (x600) shows degenerate Purkinje cells in the cerebellar cortex with spheroids (white arrows) in the B-APP immunostain (H, x400). (I) Macroscope (x4) image of the medulla showing hypoplastic pyramidal tracts. (J) Photomicrograph (x100) showing vacuolation of the pyramidal tracts.

### Whole Exome Sequencing and Cohort Resequencing

In an effort to identify additional causative genes for Martsolf syndrome we used whole-exome sequencing (WES) of affected individuals from consanguineous multiplex families (Family 4911 VI:3; Family 5196 III:3) and their healthy parents (4911 V:1 & 4911 V:2; 5196 II:1 & 5196 II:2) (**Figure 1**). The alignment, read depth and estimated heterozygous SNP detection sensitivity of each individual is given in **Table S1**[19].
Sequence variants were filtered using minor allele frequency (MAF) of < 0.001, plausibly deleterious consequence and bi-allelic inheritance.

In 4911 VI:3, ten rare homozygous variants (**Table 3**) were reviewed by manual assessment of read quality using IGV2.3 software[20] and by Sanger sequencing of selected variants for their segregation in the family. Of these only a nonsense mutation, c.452G>A, p.Trp151* (rs200086262) in *ITPA* (NM_033453, MIM 147520) segregated within the family in a manner consistent for an autosomal recessive disease-causing mutation (**Figure 1A**). This variant has been previously identified as disease associated[17] and is present in gnomAD (genome Aggregation Database) with a minor allele frequency of 0.0058%. In the affected individual from Family 5196 (III:3), homozygosity for an apparently unique 40bp deletion spanning the splice donor site of *ITPA* exon 7 was detected on WES. Subsequent Sanger sequencing confirmed this to be chr20 hg19 g.3202531-3202570del; c.456_488+7del. This deletion is likely to have been microhomology-mediated as a nine base pair perfect repeat is present at the 5’ end of the deleted region and the genomic region immediately 3’ to the breakpoint (**Figure 1D**). Both parents (5196 II:1 and 5196 II:2) were heterozygous for this mutation. Western blotting of lysates from lymphoblastoid cell lines (LCLs) from 5196 III:3 and her mother showed that ITPA protein was completely absent in the cells derived from the affected girl (**Figure 1E**). Sanger sequencing of *ITPA* in the remaining members of the cohort of 85 “mutation negative” families[8] revealed no further plausibly disease-associated mutations. The primers used for this analysis are given in **Table S2**.

**Table 3:** Rare homozygous variants detected in proband 4911 VI:3.

| Chromosome | Position | Reference nucleotide | Alternative nucleotide | rsID | Gene Name | Transcript ID | Peptide change | Segregation |
| --- | --- | --- | --- | --- | --- | --- | --- | --- |
| **1** | 220267581 | G | T | rs149324758 | *IARS2* | ENST00000302637 | Arg8Leu | No |
| **2** | 10192477 | G | A | rs199770737 | *KLF11* | ENST00000305883 | Arg461Gln | No |
| **9** | 114246291 | C | T |  | *KIAA0368* | ENST00000259335 | Gly177Glu | No |
| **13** | 42761244 | C | A | rs147914294 | *DGKH* | ENST00000536612 | Thr397Asn | No |
| **17** | 56783857 | G | A |  | *RAD51C* | ENST00000425173 | Cys170Tyr | No |
| **18** | 5478295 | C | A | rs117900256 | *EPB41L3* | ENST00000341928 | Ser109Ile | No |
| **19** | 1918914 | C | T |  | *SCAMP4* | ENST00000316097 | Ala107Val | No |
| **19** | 35524607 | G | A | rs72558029 | *SCN1B* | ENST00000262631 | Val138Ile | No |
| **20** | 3202527 | G | A | rs200086262 | *ITPA* | ENST00000380113 | Trp151* | Yes |
| **22** | 51039276 | T | G |  | *MAPK8IP2* | ENST00000329492 | Leu10Arg | No |

### Inosine ribobases (rI) are incorporated in RNA from an affected individual

*ITPA* encodes inosine triphosphate pyrophosphatase (ITPase) which hydrolyzes both inosine triphosphate (rI) and deoxyinosine triphosphate (dI)[21, 22]. Its major function is thought to be to ensure the exclusion of these “non-canonical” purines from RNA and DNA in order to avoid transcript and genome instability. Complete deficiency of ITPase in all tissues would thus be predicted to result in an increase in the incorporation of rI and dI into RNA and DNA respectively. To test this we first purified cellular RNA from a lymphoblastoid cell line (LCLs) from 5196 III:3. This RNA was digested to single nucleotides and analysed using a combination of HPLC and mass spectrometry (HPLC/MS). Using this approach we found that rI was present in RNA at a level of 725±158 SEM nucleotides of rI per 10^6^ nucleotides of AMP in 5196 III:3, a significantly higher level than in RNA from LCLs derived from either her father (17±11 SEM rI:rA x 10^6^) or her mother (71±60 SEM rI:rA x 10^6^) (**Figure 3A**). This equates to approximately one rI base in every 5500 bases of RNA from the null LCL.

**Figure 3.**
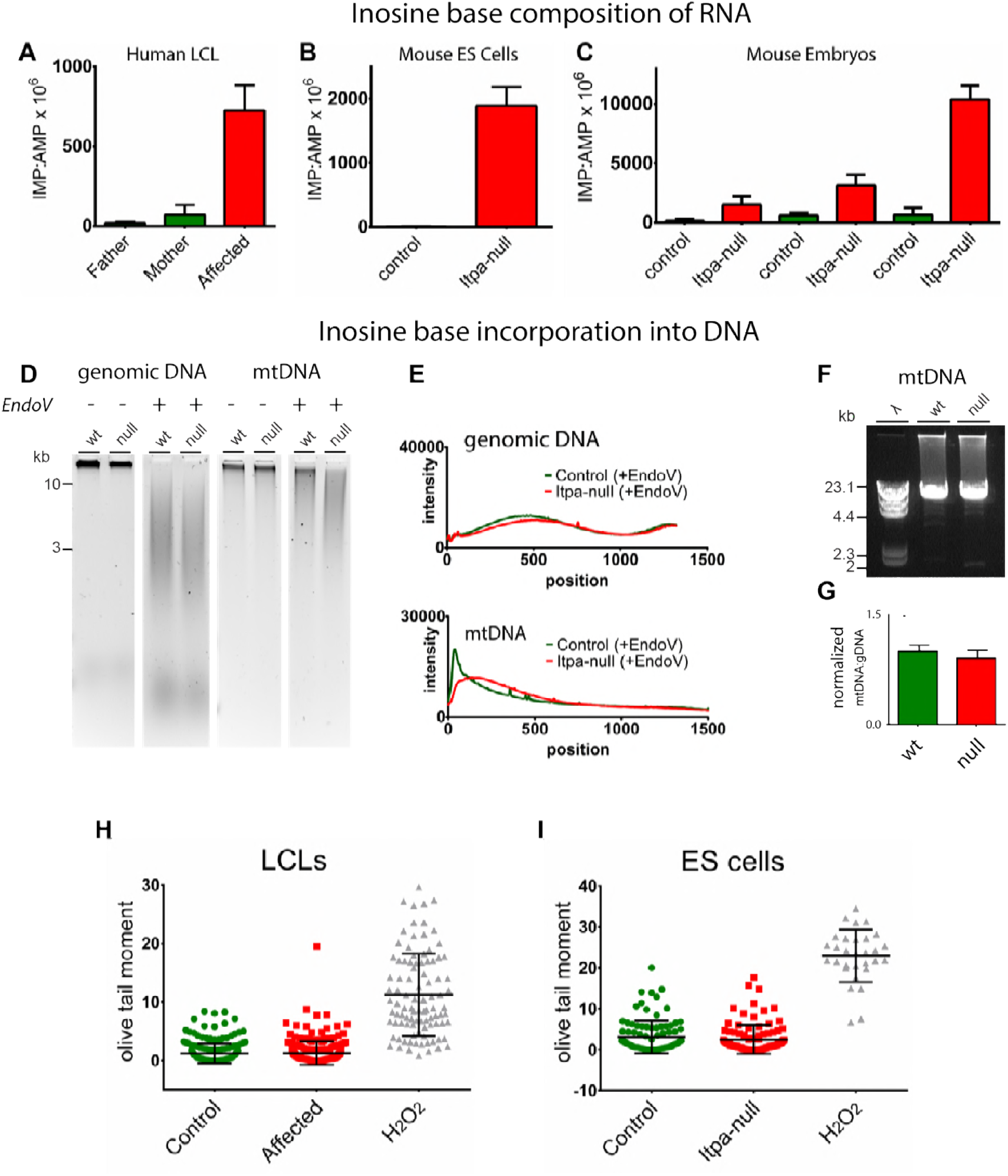
Inosine incorporation into nucleic acids in human and mouse cells lacking functional ITPase. **(A)** Bar chart showing increased inosine base content of RNA in lymphoblastoid cell lines (LCLs) derived from an affected individual (5196 III:3) as compared to those derived from her parents (5196 II:1 & II:2) **(B)** Bar chart showing increased inosine base content of RNA in *Itpa*-null mouse embryonic stem (ES) cells as compared to control ES cells. **(C)** Bar chart showing increased inosine base content of RNA derived from *Itpa*-null tissue as compared to controls. Inosine content is significantly higher in RNA derived from *Itpa*-null hearts than that derived from *Itpa*-null kidneys. Error bars ±SEM. **(D)** Alkaline-gel electrophoresis of total DNA and mtDNA extracted from mouse ES cells untreated or treated with bacterial endonuclease V (EndoV). All lanes shown are on the same gel, and these data are representative of three independent experiments. **(E)** Densitometry of gels shown in D does not identify any difference between control (green lines) and Itpa-null (red lines) cells for genomic DNA (top panel) but for mtDNA (bottom panel) there is a shift in the migration pattern in the *Itpa*-null cells suggested increase EndoV digestion compared to the controls. **(F)** Long-range PCR (LR-PCR) of the mitochondrial genome shows no evidence for increased deletions in *Itpa*-null ES cells as compared to controls. The data shown are representative of three independent experiments. The primers used are listed in **Supplementary Table S2**. **(G)** Quantitative RT-PCR (qPCR) on total DNA shows that ratios of mtDNA to genomic DNA are comparable between control and *Itpa*-null cells. The data shown are derived from analysis of six individual DNA preparations per genotype, each analysed in triplicate. All the primers used are listed in **Supplementary Table S2**. **(H,I)** Alkaline comet assays on LCLs derived from an affected individual (5196 III:3) and her mother (5196 II:2) and null and parental mouse ESC respectively with cells exposed to hydrogen peroxide as a positive control. Neither cell type shows evidence for increase single or double strand breaks in genomic DNA. Quantitation of DNA damage is by Olive tail moment (the product of the tail length and the fraction of total DNA in the tail) and is a measure of both the extent of DNA fragmentation and size of fragmented DNA.

### Inosine ribobases (rI) are incorporated in RNA from *Itpa-null* mouse ES cells and embryonic tissues

We generated *Itpa*-null mouse embryonic stem (ES) cells using CRISPR/Cas9 genome editing [23, 24] (**Figure 4A**, primers encoding the guide RNAs are detailed in **Table S2**). In these cells, rI was detectable in RNA at 1889 nucleotides rI per 10^6^ nucleotides AMP ± 295 SEM (**Figure 3B**).

**Figure 4.**
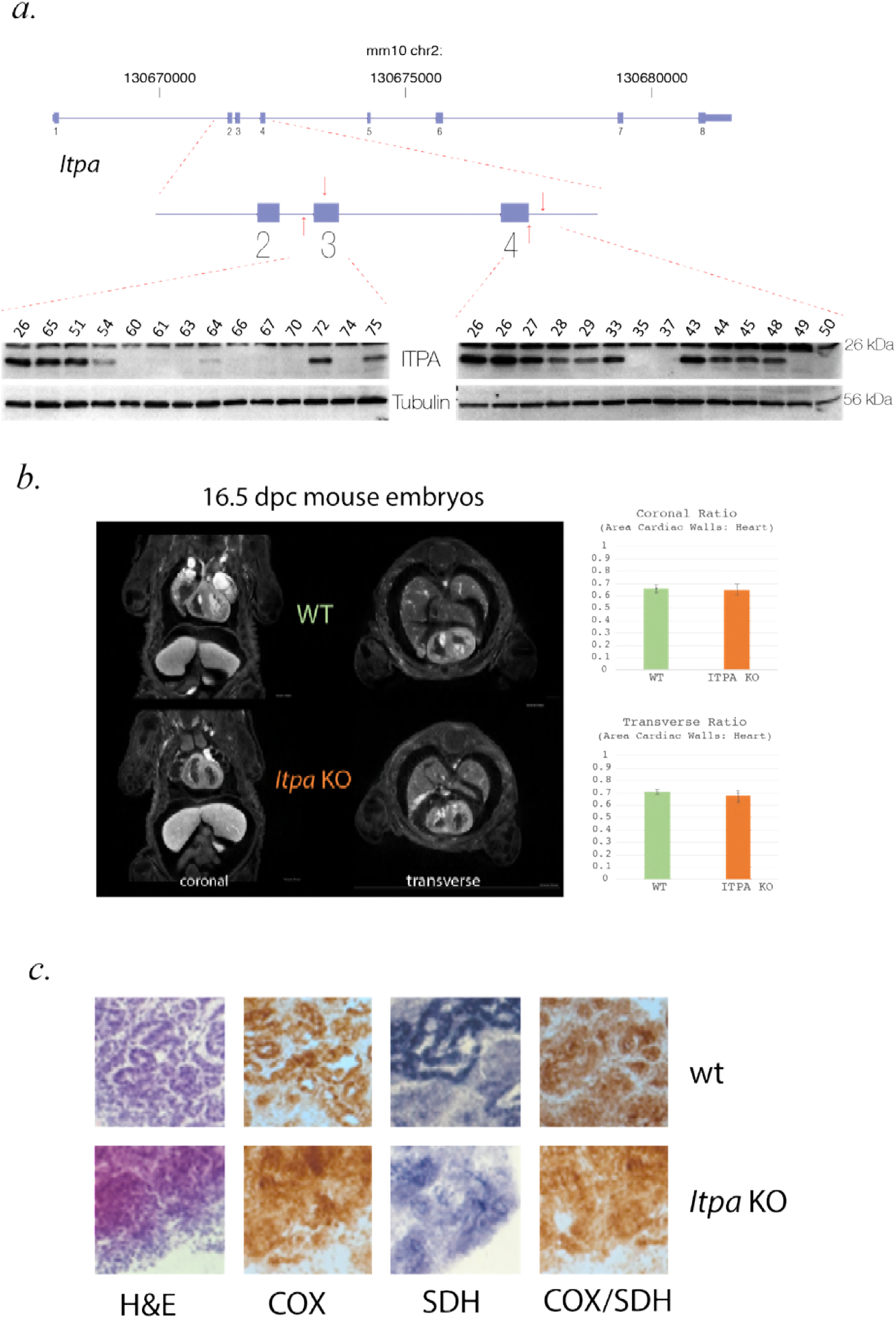
Creation, morphology and biochemical analysis of *Itpa* null mouse embryos. **(A)** Cartoon representation of the mouse *Itpa* genomic locus and gene structure with a more detailed diagram of exons 2-4 indicating the position of the guideRNAs used to create the null alleles in the mouse lines to create null embryos. Representative western blots are shown of embryonic tissue demonstrating absence of Itpa protein in samples used as “Itpa null”. ITPA protein is detected in lysates from control but not *Itpa*-null cells upon probing the blot with polyclonal antibodies raised to full-length ITPA (Millipore) and an N-terminal domain of the protein encoded by sequence 5’ of that mutated by CRISPR (LSBio). Blotting for Tubulin serves as a loading control and each lane on the blot corresponds to an individual lysate sample. **(B)** Representative coronal and transverse images through the heart from optical projection tomography (OPT) of wild-type (top panel) and *Itpa*-null (bottom panel) e16.5 embryos. The bar charts to the right of this image shows quantification of the heart wall to total heart area ratio which showed no difference between null (orange) and control (green) embryos. **(C)** Oxidative enzyme histochemistry of wild-type and *Itpa*-null embryonic heart. Sections were subjected to H&E staining, individual COX and SDH reactions together with sequential COX/SDH histochemistry. No evidence of morphological changes or focal enzyme deficiency in the Itpa-null heart was identified. Data are representative of duplicate experiments.

To determine if there was a correlation between the level of rI in different tissues *in vivo* and the organs affected in the human disease, we generated mice heterozygous for *Itpa* null alleles using direct cytoplasmic injection of Cas9 mRNA and guide RNAs into zygotes. Heterozygous animals were crossed to generate *Itpa*-null embryos and wild-type littermate embryos. Both genotyping and Western blot analysis (**Figure 4A**) were used to confirm the null status of each embryo (**Figure 4A**). As in previously reported targeted inactivation of *Itpa*, we found reduced body size (with a proportionate reduction in heart size) in *Itpa-null* embryos[25] and no other obvious morphological differences compared to wildtype controls (**Figure 4B**). The level of rI in RNA from Itpa-null hearts (10382 nucleotides IMP per 10^6^ nucleotides AMP ± 2008 SD) was significantly higher than in either *Itpa*-null brain or kidney (p<0.05 and p<0.01 respectively, student’s t-test) and equated to approximately one rI for every 385 bases of RNA (**Figure 3C**). rI was present at very low levels in RNA derived from control tissues.

### Inosine base incorporation is detectable in mtDNA but not genomic DNA

The bacterial endonuclease V (nfi/EndoV) cleaves DNA at dI bases creating nicks in the dsDNA[26]. Digestion of genomic DNA from control and *Itpa*-null ES cell lines using EndoV (New England Biolabs) followed by alkaline-gel electrophoresis revealed no measurable difference in migration between the samples (**Figure 3D**). However, a small but reproducible increase in the EndoV sensitivity of mtDNA from *Itpa*-null ES cells as compared to that in controls was seen (**Figure 3E**).

To assess whether this increased inosine incorporation was associated with increased instability of the mitochondrial genome (mtDNA), we used quantitative PCR (qPCR) to compare levels of mtDNA to levels of nuclear DNA. We also used long-range PCR (LR-PCR) of mtDNA to look for any increase in the frequency of mtDNA rearrangements. Neither assay showed any differences between *Itpa*-null cells and controls (**Figure 3F,G**), or between Itpa-null tissues and controls (**Figure S1A-B**). We used Ion Torrent sequencing to detect base substitutions and MinION sequencing to detect large-scale mtDNA rearrangements amplified from control and *Itpa*-null kidneys and hearts. No differences between wild type and *Itpa*-null kidney or heart were detected (**Figure S1C-D**).

Using HPLC/MS analyses we were unable to detect dI in the digested DNA samples from 5196 III:3, 5196 II:2 or the *Itpa*-null mouse ES cells (data not shown). To assess secondary effects of low-level dI incorporation on genome stability, a commercial comet assay kit (Trevigen) was used. No increase in DNA strand breaks could be detected in 5196 III:3 compared to 5196 II:2 LCLs, or in *Itpa*-null compared to wild-type ES cells (**Figure 3H,I**).

### Biochemical Assessment of Mitochondrial Function

To assess whether ITPase deficiency had any effect on mitochondrial function, we carried out metabolic tracer analysis on the ES cells using ^13^C_5_-glutamine, and conducted functional histopathology on tissue samples. In the tracer experiments, ^13^C-incorporation into fumarate and citrate was analysed (Figure S1E). The (m+4) isotopologues of both metabolites indicated that ITPA-loss did not affect normal oxidative TCA cycle function and a low level of reductive carboxylation of oxoglutarate, as measured by citrate (m+5), was again minimally altered upon loss of ITPA. Functional histopathology on the tissue samples was adapted from analyses used in a clinical diagnostic setting. Samples were reacted for cytochrome *c* oxidase (COX) and succinate dehydrogenase (SDH) activities, with sequential COX-SDH histocytochemical analyses carried out in order to identify low level, focal COX-deficiency. No differences were seen between control and *Itpa*-null tissues and no COX-deficient cells were identified in *Itpa*-null heart (**Figure 4C**).

### Transcriptome and Proteome Analysis of *Itpa* null cells/tissues

We compared the transcriptome of control and *Itpa*-null mouse hearts using the Affymetrix MTA1.0 expression microarray. There was very strong concordance between transcript levels in control and *Itpa*-null samples when all loci were examined together (**Figure 5A,B**) or when a subset of loci that have been implicated in dilated cardiomyopathy in mice or humans were examined separately (**Figure 5E**). When specific cardiac disease genes were examined using ddPCR, modest reductions could be observed in *Itpa*-null heart tissue (**Figure 5G**). However it was not possible to determine if reductions of this magnitude would significantly alter cardiomyocyte function and, more importantly, we could not distinguish whether these changes in transcript levels were primary effects or secondary to an early disease process in heart.

**Figure 5.**
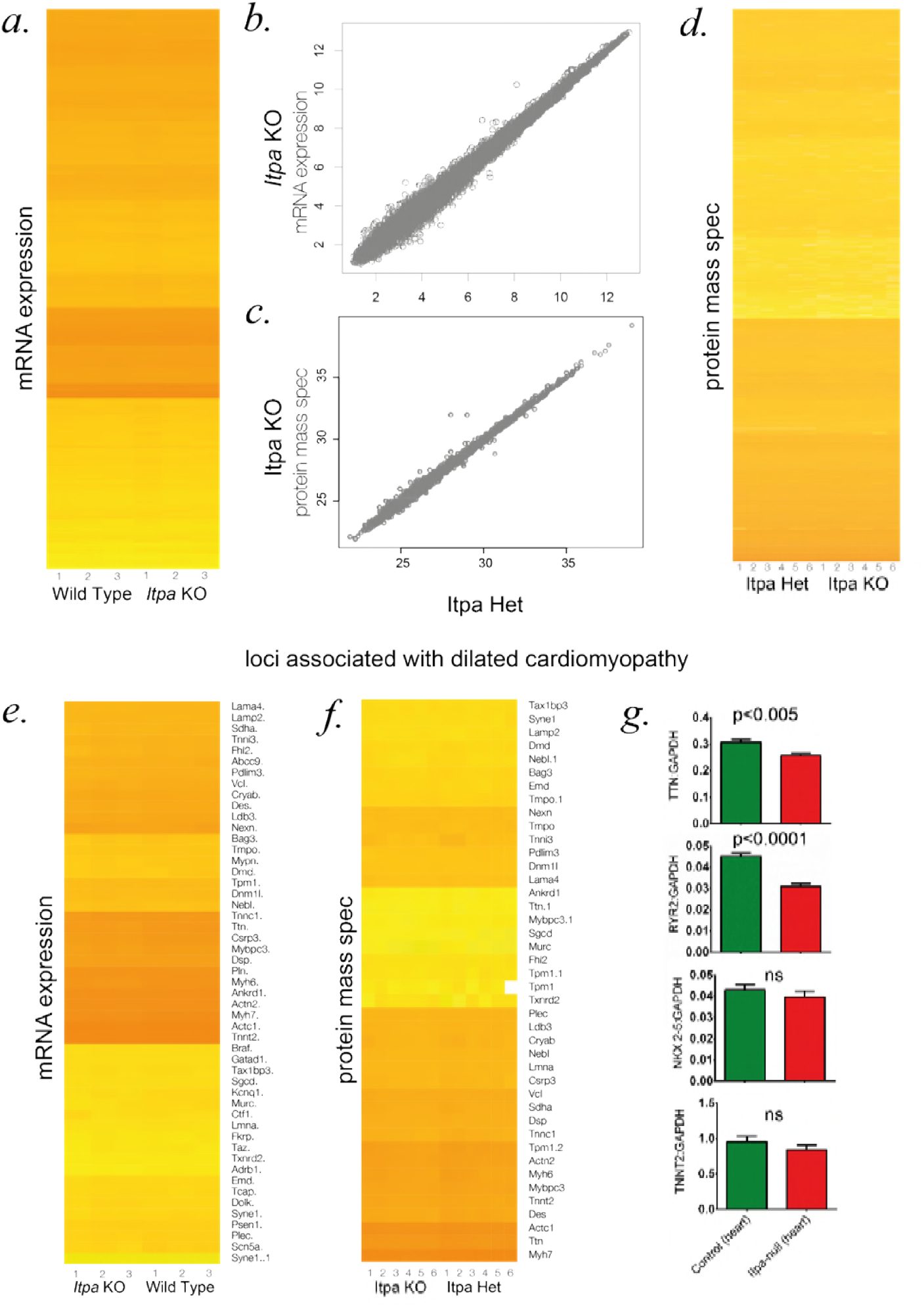
Transcriptomic and proteomic analyses of *Itpa* null heart. **(A)** Plot of per-gene log_2_ signal from Affymetrix MTA1.0 microarray on samples from wild-type and *Itpa*-null embryonic hearts. Three samples per genotype were analysed. **(B)** Heatmap.2 based clustering of per-transcript genome wide log_2_ ratios from biological triplicates of RNAseq from *Itpa*-null embryonic hearts and littermate controls. **(C)** Heatmap.2 based clustering of genome wide per-protein intensity from six biological replicates of quantitative mass spectrometry from *Itpa*-null embryonic hearts and littermate controls. **(D, E)** Subset of data from B & C focussed on transcripts and proteins that are known to be involved in mendialian causes of dilated cardiomyopathy. No major differences are detectable in any of the heatmaps. **(F)** Quantitative RT-PCR (qPCR) of selected transcripts in *Itpa*-null embryonic hearts and littermate controls. The data shown are derived from analysis of 10 individual cDNA preparations per genotype, each analysed in triplicate. All the primers used are listed in **Supplementary Table S2**.

A generalized effect on transcription produced by increased inosine incorporation into RNA in *Itpa*-null cells might not be identified on transcriptome analysis if its effects on individual transcripts are proportionate. Therefore, in order to assess any changes in transcription rate or transcript stability, we labelled RNA transcribed over the course of 30 minutes with the ribonucleotide analogue 4-thiouridine (4sU). 4sU incorporation was assayed immediately after labelling, providing a measure of transcription rate, and then at subsequent time points, at which any changes in the rate of RNA turnover would be revealed (**Figure S2A**). 4sU incorporation was assayed by biotinylation of its thiol group and then quantification of biotin using a fluorescence-based kit. When RNA was harvested immediately following treatment, 4sU incorporation appeared lower in *Itpa*-null cells than in control cells. However, this effect was not confirmed when 4sU in digested RNA was measured directly by HPLC (data not shown). Similarly, reduced incorporation was not seen when an alternative assay utilizing 5-Ethynyl Uridine was used (**Figure S2C**). No differences in the 4sU content of control and *Itpa*-null cells were observed at the later time points, suggesting that inosine incorporation does not affect RNA stability globally (**Figure S2A-B**).

Label-free quantitative mass spectrometry was performed to examine the whole proteome in control and Itpa-null mouse heart tissue. Apart from the absence of ITPase, no significant differences were detectable on inspection of either the whole dataset or the dilated cardiomyopathy-associated proteins specifically (**Figure 5C,E**).

## Discussion

Bi-allelic loss-of-function mutations were recently reported in *ITPA* in seven affected individuals from four families with early-infantile encephalopathy, a distinctive pattern of white matter disease evident on brain MR imaging, microcephaly and progressive neurological disease[17]. While no measurement of rI/dI incorporation into RNA or DNA was presented from these cases, the clinical and genetic evidence for causation was compelling in this group of children. Here we have shown that Martsolf syndrome with iDCM, is an allelic disorder. There is also evidence of phenotypic overlap between the disorders as one of the seven affected individuals reported by Kevelam et al.[17] had iDCM and three had early onset cataracts. Taken together with the existing mouse genetic data [25, 27], these data strongly support an essential role for ITPase activity in development and maintenance of brain, eye and heart function in mammals.

Since 2015 there have been no further reports of severe ITPase deficiency. The severity, the distinctive phenotype and the increasing use of whole exome sequencing in clinical diagnostics make it unlikely that this would be missed. This suggests that ITPase deficiency is genuinely very rare. In gnomAD there are 48 individuals are heterozygous for 19 different variants ultra-rare loss-of-function *ITPA* alleles. These variants have a combined MAF of 0.00024 suggesting a carrier frequency of ~1:2080 for LOF alleles in *ITPA.* Under the assumption of random mating this would give an expected birth incidence of ~1:17 million for biallelic LOF alleles in *ITPA*, which presumably explains why both families we have identified are consanguineous.

There were obvious candidate mechanisms for a cellulopathy associated with ITPase null state. First, instability of the nuclear genome induced by dITP incorporation into DNA (as seen in E coli[26]); second, instability of the mitochondrial genome via the same mechanism; third, inhibition of RNA polymerase II by rI (previously demonstrated *in vitro[28]);* fourth, instability of mature transcripts through EndoV-mediated degradation of rI-enriched mRNA; finally, induction of energy deficiency state due to biochemical perturbation of mitochondrial function. In this paper we have attempted to address each of these and failed to observe any single major effect. The differential incorporation of inosine bases between DNA and RNA is interesting. This may reflect the evolution of efficient DNA surveillance and repair mechanisms to deal with deamination of adenosine bases to form dI with the steady state for inosine in DNA being ~1 per 10^6^ nucleotides[29]. This would suggest that even moderate increases in the incorporation of dITP into DNA in ITPase null cells are likely to be below the limit of detection for the assays used here.

In this regard it is significant that we could detect low-level dI incorporation into the mitochondrial genome. Importantly we could detect no effect on either the quantity or structural integrity of mtDNA from hearts from *Itpa*-null embryos as compared to controls (**Figure S1**). The lack of evidence for a DNA-based mechanism taken together with the correlation of rI incorporation in RNA with organ severity suggested to us that there may be a transcriptomic mechanism of disease.

Enzymatic A-to-I editing in RNA is used to “recode” specific transcripts in a highly regulated manner[30] and this may explain why there is no rI-induced repair system for RNA. However, human EndoV[29] is capable of cleaving RNA at inosine bases[31, 32] and over-incorporation of rI could lead to a generalized instability of the transcriptome. We found evidence of a reduction in the transcript abundance of some longer mRNA extracted from *Itpa*-null mouse embryonic heart. This effect is difficult to interpret given that it is a relatively minor change and is plausibly a secondary effect of the disease process rather than due to rI-induced RNA instability. One way to address this problem would be to create animals who are null for both *Itpa* and *Endov* and thus determine if loss of the ribonuclease activity would rescue any or all of the *Itpa* phenotypes. Although we have not reported the details here, our preliminary work using *Itpa/EndoV* double KO mouse embryos suggested no reduction in rI incorporation into RNA.

It seems probable that the major disease mechanism in severe ITPase deficiency related to either inosine base production or rI incorporation. It is not clear why the heart, brain and developing eye are more sensitive to the perturbation. A clear understanding of the disease mechanism is important as it may lead to therapies that will ameliorate the progressive cardiac and neurological effects of this rare but important disease.

## Methods and Models

### Clinical samples and information

Our cohort consists of DNA samples from 85 families, submitted by referring clinicians for research screening for Warburg Micro syndrome and Martsolf syndrome[8]. Affected individuals in this cohort are negative for causative variants in the coding sequences of *RAB3GAP1, RAB3GAP2, RAB18* and *TBC1D20*, the genes previously associated with these disorders. Informed consent has been obtained from all participating families, and studies were approved by the Scottish Multicentre Research Ethics Committee (04:MRE00/19).

### Whole exome sequencing and analysis

DNAs from Families 4911 and 5196 (nuclear trios) were enriched for exonic sequence using kits indicated in **Table S1** and sequenced using Illumina HiSeq technology as described previously[33]. Sequence reads were aligned to the GRCh37 human genome reference assembly with BWA mem 0.7.10[34]. Duplicate reads were marked with Picard MarkDuplicates 1.126. Reads were re-aligned around indels and base quality scores recalibrated with GATK 3.3[35]. Single nucleotide variants and small indels were called with GATK 3.3 HaplotypeCaller on each sample and GenotypeGVCFs to produce a raw variant call set. Variants were annotated using the Ensembl Variant Effect Predictor[36]. Statistics for alignment, read depth and estimated heterozygous SNP detection sensitivity for each individual are listed in **Table S1**. Sequence variants were filtered using minor allele frequency (MAF) of < 0.001, plausibly deleterious consequence and bi-allelic inheritance.

### Sanger sequencing

PCR amplification of the coding exons of *ITPA* and intron-exon boundaries was carried out using flanking primers with M13 tags to facilitate later sequencing (see **Supplementary Table S2**). Primers were designed using ExonPrimer software on the basis of the reference sequence NM_033453. Sequencing reactions were carried out with BigDye Terminator 3.1 reagents (Applied Biosystems), according to manufacturer’s instructions. Sequencing data was analysed with Mutation Surveyor software (SoftGenetics).

### Cell culture

Cells were maintained under 5%CO_2_ at 37°C. Lymphoblastoid cell lines (LCLs) were maintained in suspension in RPMI1640 media (Gibco) supplemented with 10% foetal calf serum, 1mM oxalocacetate, 0.45mM pyruvate, 0.03% glutamine, 1% penicillin/streptomycin and 0.2 I.U/ml insulin. Embryonic stem (ES) cells were maintained in adherent culture in GMEM supplemented with 10% foetal calf serum, 0.1mM non-essential amino acids, 2mM L-Glutamine, 1mM sodium pyruvate and 106 units/L LIF.

### Antibodies and Western blotting

Rabbit polyclonal antibody raised to full-length ITPA was obtained from Millipore. Rabbit polyclonal antibody raised to an N-terminal portion of ITPA was obtained from LSBio. Goat polyclonal antibody raised to β-tubulin was obtained from Abcam.

Cells were lysed in a buffer containing 0.5% (v/v) Nonidet P-40 in a solution of 150mM NaCl, 10mM EDTA and 50mM Tris-HCl (pH=7.5) to which a protease inhibitor cocktail (Roche) was added. Tissue samples were lysed directly in 1x NuPage LDS Sample Buffer (Thermo Fisher) containing 5% β-mercaptoethanol. SDS-PAGE and Western blotting were carried out according to standard methods. ECL 2 Western blotting substrate (Pierce) was used to produce chemiluminescent signal, HyperFilm ECL (General Electric) was developed using a Konica Minolta SRX-101A.

### Detection of Inosine bases in hydrolysed RNA and DNA

Cellular RNA and DNA were purified using RNAeasy (Qiagen) and BACC2 (GE Healthcare) kits respectively. mtDNA isolation was performed using a mitochondrial DNA isolation kit (Abcam).

For analysis of nucleic acid composition by mass spectrometry, digestion to single nucleotides was carried out. A combination of either 50μg/ml RNAseA (RNA) or 20U/ml DNAseI (DNA)(Roche Diagnostics) respectively and 80U/ml NucleaseP_1_ (Sigma) was used as previously described[37]. Both purification and digestion was carried out in the presence of a 20μM concentration of the adenosine deaminase (ADA) inhibitor deoxycoformycin (DCF)(Sigma). Digestions were carried out in a buffer containing 1.8mM ZnCl_2_ and 16mM NaOAc, pH=6.8 at 37°C overnight. Nucleases were then removed with 10,000 MW cut-off spin columns (Amicon). Samples were loaded onto a ZIC-pHILIC column using a Dionex RSLCnano HPLC and the eluate was applied to a Q Exactive mass spectrometer in negative mode. The instrument was operated in tSIM mode and data were quantified using XCalibur 2.0 software.

For analysis of genomic and mitochondrial DNA composition by Endov-digestion and alkaline-gel electrophoresis, DNA samples were treated with 10 U of Endonuclease V (NEB) with the supplied buffer for 2 hours at 37°C. DNA strands were separated by incubation at 55°C in loading buffer containing 3% Ficoll (type 400) and 300mM NaOH. Samples were separated on agarose gels (50mM NaOH, 1mM EDTA) with a solution of 50mM NaOH, 1mM EDTA used as running buffer. After electrophoresis, gels were neutralized and stained with SYBR Gold (Invitrogen).

### Comet assays

Alkaline comet assays were carried out using the Trevigen CometAssay electrophoresis kit according to manufacturer’s instructions. Briefly, cells were embedded into low melting agarose on comet slides and incubated in lysis solution overnight in the dark at 4°C. They were incubated in a solution of 300 mM NaOH, 1 mM EDTA for 30 minutes at room temperature, then electrophoresed in this solution for 30 min at 21 volts at 4°C in the dark. Comet slides were immersed in 70% ethanol for 10 min at room temperature and dried at 37°C for 15 minutes. They were then stained with 1x SYBR gold in TE buffer (pH 7.5) for 30 minutes at room temperature, dried for an additional 15 minutes at 37°C and visualized with a Zeiss Axioskop 2 epifluorescence microscope with a 10x objective. Data were analysed with CaspLab 1.2.3 software.

### Long-range PCR of mitochondrial DNA

10ng DNA samples were amplified using the TaKaRa LA Taq polymerase mix with primers flanking nucleotide positions 272 and 16283 on the mouse mitochondrial genome (NC_005089; see **Supplementary Table S2**). A long PCR template program was used as follows: 94°C – 2 minutes, 35 cycles of 94°C – 30 seconds and 65°C – 16 minutes followed by a final extension of 72°C – 16 minutes.

### Quantitative PCR

For quantitative PCR (qPCR) of mitochondrial (mtDNA) and genomic DNA (gDNA), DNA preparations (retaining both species) were made from cell and tissue samples using Viagen reagent (Viagen Biotech) according to manufacturer’s instructions. For analysis of gene expression by qPCR, RNA was extracted using Trizol reagent together with an RNeasy mini kit (Qiagen) according to manufacturer’s instructions. Purified RNA was used immediately as a template for production of cDNA using a First Strand cDNA Synthesis Kit for RT-PCR (AMV) (Roche).

qPCR analysis was carried out on a LightCycler 480 (Roche). Amplification from mtDNA and gDNA was carried out using pairs of primers designed to amplify from the mtCO1 locus of the mitochondrial genome and the *Gapdh* locus of the autosomal genome. Amplification from cDNA was carried out using primers designed to amplify from TTN, RYR2, TNNT2 and GAPDH cDNAs. Amplification from NKX2-5 was carried out using commercial TaqMan probes (Mm01309813_s1_Nkx2–5). PCR amplification with unlabelled primers was quantified through binding of specific mono color hydrolysis probes (Roche). Data were analyzed using LightCycler 480 software version 1.5.0 (SP4) (Roche). Primers were designed using the Universal ProbeLibrary Assay Design Center and are listed in **Supplementary Table S2**.

Droplet Digital PCR (ddPCR) reactions were carried out according to manufacturer’s instructions (Biorad). In each reaction, cDNAs were combined with a VIC-labeled TaqMan control probe, mouse GAPD or eukaryotic 18S (Life Technologies). Primers specific for target genes are as above. Droplets were generated using a Biorad QX200 or QX200AutoDG droplet generator, PCRs were carried out using a C1000 Touch Thermal Cycler, and droplets were analyzed on a QX100 Droplet Reader. The data were analyzed using Quantasoft software (QuantaLife).

### Mitochondrial enzyme histochemistry

Tissue samples from Itpa-null embryos and littermate controls (e16.5-e18.5) were frozen in liquid nitrogen-cooled isopentane prior to sectioning. Cryostat sections were stained for individual activities of COX and SDH and also for sequential COX/SDH activity. Briefly, sections were reacted for 45 min at 37 °C with COX reaction media (4 mM diaminobenzidine tetrahydrochloride, 100 μM, cytochrome *c* and 20 μg/ml catalase in 0.2 M phosphate buffer, pH 7.0) and 40 min at 37 °C with SDH media (1.5 mM nitroblue tetrazolium, 1 mM sodium azide, 200 μM phenazine methosulphate, 130 mM sodium succinate, in 0.2 M phosphate buffer, pH 7.0).

### Microarray

RNA was extracted from e16.5 mouse hearts using Trizol reagent together with an RNeasy mini kit (Qiagen) according to manufacturer’s instructions. RNA quality was assessed using an Agilent Bioanalyser instrument and Total RNA nano. RNA integrity numbers (RIN) were ≥9.1 for all samples. Transcriptome analysis was carried out by Aros Applied Biotechnology A/S using the Affymetrix MTA1.0 microarray. Data were analysed using Affymetrix Transcriptome Analysis Console 3.0 and custom R scripts

### Label-free quantitative proteomics

Protein was extracted from embryonic mouse hearts in a buffer containing 8M Urea, 75mM NaCl and 50mM Tris, pH=8.4 by sonication at 0-4°C in a Bioruptor device (Diagenode) together with silica beads. Protein concentrations were quantified using a BCA assay (Pierce) and then 100μg of each sample was subjected to in-solution tryptic digest. Samples were loaded onto a C18 column using a Dionex RSLC Nano HPLC and the eluate was applied to a Q Exactive mass spectrometer. The data were quantified using XCalibur 2.0 software.

### Generation of Itpa-null mouse ES cells and embryos

*Itpa*-null mouse ES cells were generated using CRISPR/Cas9 genome editing [23, 24]. Paired guide RNA (gRNA) sequences were selected using the online CRISPR design tool at(http://crispr.mit.edu/). Oligonucleotides encoding these sequences (**Supplementary Table S2**) were annealed and ligated into p✗461 and p✗462 plasmids (Addgene). Recombinant plasmids were verified by direct sequencing. For each targeted locus, the E14 ES cells were transduced with 1μg of each vector using the Neon system (Life Technologies) according to manufacturer’s instructions. Cells were allowed to recover for 24 hours, then treated for 24h with 1 μg ml^-1^ puromycin in order to select for cells containing the px462 construct. To select single cells also containing the px461 construct, fluorescence activated cell sorting into 96-well plates was carried out using a FACSJazz instrument (BD Biosciences). Clonal cell lines were analysed by direct sequencing of targeted alleles and by Western blotting. Sequencing primers are shown in **Supplementary Table S2**. To facilitate sequence analysis, PCR products were cloned into pENTR/D-TOPO vectors prior to sequencing.

Cytoplasmic zygotic injection of wild-type Cas9 mRNA together with *in vitro* transcribed gRNAs was used to generate Itpa-null mouse embryos. This approach was also used to produce heterozygous-null animals used to establish transgenic mouse lines. The plasmid vectors described above were used as a template for PCR amplification together with forward primers incorporating T7 promoter sequences and a universal reverse primer (see **Supplementary Table S2**). RNA was synthesised using a HiScribe T7 High Yield RNA Synthesis Kit (New England Biolabs) according to manufacturer’s instructions. DNA preparations were made from tissue samples using Viagen reagent (Viagen Biotech). Genotyping was carried out by direct sequencing of targeted alleles and by Western blotting as above. Following initial genotyping of Itpa-null animals produced by crossing heterozygous-null founders, subsequent genotyping of transgenic lines was conducted through PCR analysis.

### Optical projection tomography and morphometry

E16.5 mouse embryos were mounted in 1% agarose, dehydrated in methanol and then cleared overnight in a solution containing 1 part Benzyl Alcohol and 2 parts Benzyl Benzoate. Imaging was conducted with a Bioptonics OPT Scanner 3001 (Bioptonics, UK) using brightfield analysis to detect tissue autofluorescence for capture of anatomical and signal data (wavelengths: excitation at 425 nm, emission: 475 nm). The resulting data were reconstructed using Bioptonics proprietary software (Bioptonics, MRC Technology, Edinburgh, UK), automatically thresholded to remove background signal, then merged into a single 3D image output using Bioptonics Viewer software. Measurements of internal chest cavity diameter, maximum heart diameter, cardiac wall cross-sectional area and total heart cross-sectional area were taken for five embryos per genotype.

## Supplemental Data

Supplementary data consists of two tables and three figures.

## Acknowledgements

We would like to thank the WARBM children and their families.
http://warburgmicrosyndrome.org/. DRF, AJP, MST and IA were each supported by program grants to Medical Research Council Human Genetics Unit. MTH is supported by a research grant from the Newlife Foundation for Disabled Children (R43152; 13–14/02). RWT is supported by the Wellcome Centre for Mitochondrial Research (203105/Z/16/Z), the Medical Research Council (MRC) Centre for Translational Research in Neuromuscular Disease, Mitochondrial Disease Patient Cohort (UK) (G0800674), the Lily Foundation and the UK NHS Highly Specialised Service for Rare Mitochondrial Disorders of Adults and Children. We thank Dr. Laura Greaves (Wellcome Centre for Mitochondrial Research, Newcastle University) for kindly providing long-range PCR primer sequences. We thank the staff at the IGMM transgenic core facility, and Craig Nicol for assistance with graphic design. We thanks Richard Clark and Lee Murphy from the Edinburgh Clinical Research Facility for assistance with MinION sequencing

## Web Resources

OMIM: http://www.omim.org/

Picard: http://picard.sourceforge.net/

EVS: http://evs.gs.washington.edu/EVS/

1000 genomes: http://www.1000genomes.org/

ExAC: http://exac.broadinstitute.org/

## Supporting Information Legend

There is a single supporting information file associated with this manuscript which contains two tables, two figures and the associated methods and references. **Table S1** documents the technical metrics relating to the trio whole exome sequencing on families 4911 and 5196. **Table S2** lists the oligonucleotide primers used for; i) sequencing the candidate human genes (including ITPA); ii) creation of genome editing reagents; iii) sequencing mouse *Itpa;* iv) mitochondrial genome analysis; v) quantitative RTPCR. **Figure S1** is titled “Sequencing of mtDNA in control and *Itpa*-null tissues and metabolic analysis of mitochondrial function in control and Itpa-null ES cells”. **Figure S2** is titled “Transcription rate and transcript stability in control and *Itpa*-null ES cells”. Legends are provided for both figures. The methods section describes the approach to sequencing using the Ion Proton and MinION, the metabolic tracer experiments and the 4-thiouridine & 5-Ethynyl Uridine incorporation assays. A related reference is provided at the end of the file.

